# VesiScope: a stand-alone tool for the automatic detection, quantification and size analysis of GUVs from phase contrast microscopy images

**DOI:** 10.64898/2026.10.08.757674

**Authors:** Dario Cecchi, Angelo Lasala, Federica Poli, Eugenia De Remigis, Sara Moccia, Stefano Palagi

## Abstract

Giant Unilamellar Vesicles (GUVs) are cell-sized phospholipid compartments that serve as versatile platforms for studying membrane-associated processes and engineering synthetic cell-like systems with applications spanning biophysics, synthetic biology, and, more recently, microrobotics. Fluorescence labeling is commonly used to enhance microscopy image contrast and facilitate automated Giant Unilamellar Vesicles (GUVs) quantification. Nonetheless, fluorescent probes may introduce unaccounted variations compared to the intended pristine formulation. Alternatively, phase-contrast microscopy enables label-free GUVs observation, but the lack of reliable automated detection methods limits high throughput analyses. To address this gap, we developed a framework based on YOLOv11 model, a state-of-art lightweight convolutional neural network for object detection. Trained on 4666 manually annotated GUVs from 164 phase-contrast microscope images, the model enables robust detection under low contrast-to-noise ratio and heterogeneous illumination. The trained model is implemented in a stand-alone desktop application supporting multiple analysis in a single run, pixel-to-micrometer calibration, and automated size distribution analysis. Retraining scripts compatible with standard hardware are also provided to enhance adaptability. VesiScope achieves high detection accuracy and competitive cross-microscope generalization enabling fast and reliable analysis of unaltered GUVs, supporting experimental reproducibility. By enabling rapid, reproducible, and label-free characterization of GUV populations, VesiScope provides a versatile tool for applications ranging from membrane biophysics and synthetic cell engineering to emerging GUV-based microrobotics, without introducing potentially perturbing fluorescent labels.

## Introduction

Giant Unilamellar Vesicles (GUVs) are micron-sized aqueous compartments enclosed by a single phospholipid bilayer, typically ranging from 1 to ∼100 μm in diameter, and are widely employed as model systems in membrane biology, synthetic biology, and bio-inspired engineering. Their cell-like dimensions and unilamellar architecture provide a simplified yet powerful platform for investigating membrane structure, dynamics, and associated biological functions. GUVs have been extensively used to study membrane biophysics[1], characterize transmembrane proteins and their interactions with lipid bilayers[2,3], and investigate the transport of molecules across biological membranes[4–6]. In addition, the possibility to integrate a wide variety of functions, either encapsulated in the aqueous lumen or embedded into/onto the membrane, has made them a central synthetic biology tool employed in the assembly of synthetic (or artificial) cells, capable of mimicking and interacting with natural cells[7–16] and, more recently, their versatility has opened new pathways in microrobotics, where GUVs can be programmed for responsive and adaptive behaviors[17,18].

Various GUV preparation methods have been established to meet different experimental needs. Emulsion-based methods are preferred for cargo encapsulation due to high encapsulation efficiency, but they can lead to traces of oil in the lipid bilayer. Swelling-based techniques are instead better suited for membrane biophysics studies, as they allow for a more precisely controlled membrane composition[19,20]. Regardless of the preparation method employed, the evaluation of GUV yield and size distribution is essential to quantify the effects of varying vesicles composition and process parameters, as well as for ensuring production efficiency and experimental reproducibility. Common strategies for characterizing GUV populations include flow cytometry[21], image-based quantification through microscopy[22,23] and a combination of both[24,25]. Fluorescent labeling is essential for flow cytometry-based analysis and facilitates automated microscopy images segmentation by enhancing the contrast between GUVs and background, thereby facilitating automated image segmentation and quantitative analysis. While providing a powerful mean to automate yield and size distribution analysis, adding fluorescent probes may unintentionally alter yield and properties of the obtained GUVs by perturbing bilayer structure and mechanics[26,27] or inner solution density[28,29].

When downstream applications of GUVs do not need fluorescent probes, direct assessment by phase-contrast optical microscopy provides a label-free strategy to quantify vesicle yield and size distribution without perturbing GUV composition. However, automated object detection in phase-contrast images is challenged by halo artifacts and low contrast-to-noise ratio (CNR), making conventional threshold-based segmentation unreliable and restricting the scalability of image-analysis workflows through the need for extensive manual intervention. As a result of intensity overlap and low contrast-to-noise ratio (CNR), conventional threshold-based segmentation methods often fail, leading to reduced detection accuracy, limited throughput, and the need for extensive manual intervention, thereby constraining the scalability of standard image-analysis pipelines[30]. To overcome these limitations, object-detection approaches based on deep learning (DL) have attracted increasing attention, owing to their robustness to low CNR and their ability to extract discriminative features even when intensity differences between objects and background are minimal[31]. To date, applications of this strategy have focused primarily on cell detection[32–34]. However, differences in morphology, refractive-index distribution, and boundary contrast prevent the direct transfer of such models to GUV detection without dedicated training, and existing AI-based approaches applied to GUV analysis remain limited to fluorescence microscopy[35,36].

To address this gap, we trained a lightweight Convolutional Neural Network (CNN) on a dataset of 164 phase-contrast images comprising 4666 manually annotated GUVs. The trained models are distributed through a stand-alone desktop application, allowing automated acquisition and processing of data from unlabelled samples, eliminating the need for fluorescent dyes that may alter GUV properties. The resulting AI-based framework enables robust high-throughput detection, quantification, and size distribution analysis of GUV populations directly from phase-contrast images. Notably, the model maintained competitive detection performance on out-of-distribution images acquired from a microscope different from that of the training, showing good generalization across imaging systems.

## Methods

### Materials and instruments

The complete list of materials necessary for GUV preparation was previously reported[37]. Glass slides were purchased from Menzel Gläser (cat. 631-0701), while cover slides and the Burker chamber were purchased from Brand (cat. 470045 and 718920). Images were acquired with two different microscopes for generalization assessment: - **microscope 1 (*μs1*):** Nikon Eclipse TE2000-U equipped with 20x and 10x objectives (cat. MRP46202 and MRP40102) and a CMOS camera (Tucsen TrueChrome PDAF paired with the 20X objective and Basler MED ace 8.9 MP, 4092 x 2160, 42 color, USB paired with the 10X objective) - **microscope 2 (*μs2*):** Leica DMi8 Inverted Epifluorescent Microscope equipped with 10x objective (cat. 506400) and CCD camera (Leica DFC7000 T).

### GUV preparation and imaging

GUVs were produced by the droplet transfer method as we previously described in [37]. Briefly, an interface was allowed to form in a 1.5 mL tube between a 240 mM glucose aqueous solution (outer solution, OS) and a DOPC in oil solution (lipid solution, LS). A 2.5% water-in-oil emulsion was prepared by vortexing a 240 mM sucrose aqueous solution (inner solution, IS) and LS, then water droplets were converted into GUVs by depositing the emulsion on top of the OS/LS interface and centrifuging the samples at 3300g for 20 minutes. The observed GUVs belonged to several different samples prepared from LS consisting of 0.2, 1 and 3.18 mM DOPC in mineral, paraffin or silicone AR20 oil. 20 μL of the resulting vesicle suspensions were deposited onto glass slides or glass-bottom Burker chambers to acquire phase contrast images and save them in RGB format as JPG files.

### Dataset curation

The acquisitions were manually annotated by six expert researchers enclosing every visible GUV in a bounding box through Label Studio[38], a widely adopted open-source image annotation tool. The dataset was divided in two main group: a set of 269 images acquired with *μs1* for model training (In-Distribution, ID) and a set of 26 images acquired with *μs2* to test cross-instruments generalization (Out-Of-Distribution, OOD). This setting was designed to probe the model’s ability to generalize across different imaging conditions, which may introduce a domain shift in optical characteristics, image appearance, and typical GUV density per image. The ID set was partitioned at the acquisition level into training, validation and test sets to prevent data leakage between splits. Leica images were reserved solely for cross-microscope generalization assessment. A detailed description of the dataset is reported in Figure S1 and in the online repository[39].

### Model Architecture, Training, and Evaluation

We used YOLOv11 (Ultralytics)[40], the latest generation of the YOLO single-stage object detection family, a lightweight architecture offering state-of-the-art performance across diverse biomedical imaging applications with reduced computational requirements for both training and deployment, in line with EU ethics guidelines on trustworthy AI [41,42].

The model performs bounding box regression and class prediction in a single forward pass and is available in multiple capacity variants, including *nano* (*n*, 2.5M parameters), *small* (*s*, 9M), and *medium* (*m*, 20M). All variants share the same backbone and detection head design, differing only in depth and width scaling factors. For each detected instance, the model returns a bounding box with a confidence score reflecting its estimated certainty that the box contains a GUV. All three variants were trained and evaluated in this study, to identify the best trade-off between detection accuracy and model size.

Because YOLOv11 requires input images with a 640×640 pixels resolution, each manually annotated image was partitioned into a grid of non-overlapping sub-images that were rescaled to the required resolution. The tiling strategy was tailored to the acquisition magnification to preserve GUVs spatial detail while maintaining architectural compatibility. Specifically, images acquired with higher magnification (20X) were split in 4, while 10X images were split in 16 tiles. This way, GUVs imaged with different magnifications are coherently represented in the input space of the model (see Tiling strategy in the online dataset repository[39]).

Training was performed with the on-the-fly data augmentation pipeline, which includes mosaic augmentation, random horizontal and vertical flips, HSV color jitter, and random scaling and translation. All models were trained for 100 epochs with a batch size of 16 and learning rate of 0.002 on an NVIDIA Tesla T4 GPU, following the standard Ultralytics training procedure with AdamW optimizer and early stopping with a patience of 50 epochs.

To assess the influence of color representation, all experiments were conducted in two parallel configurations: one using the original RGB images, and one using the very same images converted to gray scale. We chose to perform this comparison, motivated by the inherently achromatic nature of phase-contrast microscopy, to assess whether color channels carry information useful for detection or not.

Model performance was evaluated using standard object detection metrics. Precision (P) quantifies the fraction of predicted bounding boxes that correctly overlap a manually annotated box, representing the ground truth (GT); Recall (R) quantifies the fraction of GT GUVs that were successfully detected. The first is defined as the fraction of true positives (TP) over the sum of true positives and false negatives (FN), while the latter is defined as the fraction of TP over the sum of TP and false positives (FP). Two thresholds affect these values. The Intersection over Union (IoU) threshold is used to evaluate the minimum overlap between predicted and GT bounding boxes, while the confidence threshold determines how certain the model must be that an object is present before reporting a detection, thus affecting both precision and recall. A low confidence threshold results in more predicted GUVs being retained, increasing recall but also the likelihood of false positives. Conversely, a high confidence threshold filters out uncertain predictions, reducing false positives but potentially lowering recall by excluding true GUVs. The primary summary metric, mean Average Precision at IoU 0.50 (mAP50), mitigates the dependence on a specific confidence threshold by evaluating the precision–recall trade-off across the full range of confidence values. Rather than measuring performance at a single operating point, mAP50 summarizes detection accuracy over all confidence thresholds into a single score, enabling more objective comparisons between models. For the cross-microscope generalization experiments, the F1 score, the harmonic mean of precision and recall, is also reported as a single balanced measure of detection quality. To estimate variations in detection bias according to GUV diameters, we introduced the correction factor *ρ* as the ratio between Precision and Recall (for a complete explanation refer to the readme file of the online repository[43]).

## Results

### Detection performance

In the ID test set, all three model variants achieved comparable performances regardless of the images color space, with mAP50 ranging from 0.863 to 0.881. Despite the increase in the number of parameters from *nano* to *medium* model, the maximum improvement in mAP50 was only 0.011, indicating that scaling to larger variants yields marginal benefit on this task. The *nano* model trained with and tested on grayscale images attained the highest mAP50, marginally outperforming both the *small* and *medium* grayscale variants and all RGB configurations. This plateau suggests that the visual complexity of phase-contrast GUV images, where targets appear as circular, optically uniform structures against a quasi-homogeneous background, lies well within the representational capacity of the smallest YOLOv11 architecture. The results also suggest that the color channels do not carry information relevant to GUV detection by the model (S1 Table).

The results of the OOD test set revealed instead a pronounced effect of input modality. RGB models failed to transfer across images acquired with a different microscope, with mAP50 reduced more than 75% relative to the ID test set performance. By contrast, models trained on grayscale images retained higher performances (mAP50 0.514–0.525) when tested on grayscale images. These findings indicate that color channels can introduce spurious features that are uncorrelated to the physical contrast mechanism of phase-contrast microscopy, thereby reducing model generalization. Because image formation in phase-contrast microscopy is driven by refractive-index variations rather than color, removing color information improves robustness without sacrificing relevant image content. The absence of a consistent model size-related benefit was confirmed also in the OOD setting, where grayscale mAP50 values spanned only 0.011 across the three variants. Taken together, these results show that YOLOv11-*n* trained on grayscale images provides the best trade-off between accuracy and computational cost, achieving the highest in-distribution mAP50 (0.881) and competitive cross-microscope generalization (0.514) at only 2.5M parameters (S1 Table).

To evaluate whether model performance is affected by GUV diameter, we performed size-stratified evaluation of the grayscale nano model. We found out that detection performance was limited for GUVs smaller than 5 µm, improved markedly in the 5–10 µm range, and stabilized for GUVs ≥ 10 µm in diameter. The model’s highest performance coincides with the most populated range (5–15 µm), and detection quality is maintained even for the largest GUVs (> 35 µm) despite their lower abundance in the training dataset (Fig 2 C and D). Because GUV detections is biased by false positives and false negatives (decreasing precision and recall respectively), we also evaluated whether a correction factor could be introduced and whether it remained stable across different datasets and GUV sizes. We defined *ρ* as the ratio between precision and recall (see methods and the readme file of the online repository[43]) and found it to be approximately constant for GUVs ≥ 5 µm in the ID dataset over confidence thresholds ranging from 0.25 to 0.40 (Fig 2 C and S1 Fig). In contrast, *ρ* exhibited substantially greater variability in the OOD dataset (Fig 2 D) indicating that, although the model generalizes well to unseen data, reliable quantification of GUV yield and size distributions benefits from dataset-specific calibration using a representative set of manually annotated images.

### *VesiScope* Application

The trained models are distributed as part of a stand-alone easy-to-install desktop application, ***VesiScope***[43], designed following lightweight deployment principles: it requires no programming expertise, external dependencies, or a dedicated GPU, enabling local execution, thereby preserving data ownership and control (Fig 3).

The detection pipeline begins by dividing each input image into tiles using a configurable split factor. This tiling step addresses limitations on the number of objects that the model can predict per image and improves detection of small vesicles by presenting them at a scale more consistent with model training conditions. YOLO inference is then run on all tiles, and detections are recovered in the original image coordinate space. For each detected GUV, the diameter is estimated from the predicted bounding box and the size distribution is characterized by fitting a log-normal distribution to the diameter measurements using maximum likelihood estimation, yielding the log-mean (*μ*) and log-standard-deviation (*σ*) of the population.

The application supports both single-folder and batch processing modes according to the input specified in the path selection field. In batch mode, all sub-folders are first processed collectively to produce aggregate statistics, and then each subfolder is analyzed independently. An interactive calibration window (Fig 3 B) allows the user to manually specify two reference points on the first image and enter the corresponding real distance in micrometers (µm); the resulting µm/pixel conversion factor is then applied uniformly across all images in the folder, ensuring measurement reproducibility across acquisitions. Unchecking the interactive calibration option, this operation will be executed using the conversion factor (µm/px) specified by the user.

The application generates a CSV file with GUV diameters expressed in µm, a PDF report showing the size distribution histogram with fitted log-normal curve, median, and interquartile range, and a JSON file formatted for direct import into Label Studio to support manual correction, ensuring standardized and reproducible outputs across independent experimental runs. A predict folder is created containing the bounding boxes with associated confidence scores overlaid on each image crop, enabling visual inspection of model predictions as a quality control step prior to any downstream analysis or retraining, and a labels folder with a TXT file per image containing coordinates of detected bonding boxes.

### Images semi-automatic annotation and model re-training

To support continuous model generalization improvement, we developed an iterative refinement pipeline inspired by continuous training principles, which enables model adaptation to new imaging conditions without requiring *de novo* manual annotation.

As a first step, detection sensitivity on OOD data can be enhanced by lowering the confidence threshold. While this adjustment often leads to multiple detections of the same vesicle, it generally increases the overall detection rate of GUVs. The resulting JSON output, being directly import-ready for Label Studio, enables a human-in-the-loop correction workflow where users can inspect, add, or remove predicted annotations following the guidelines described in the repository *Import Images and Pre-annotation with JSON file*[43], producing a curated dataset that reflects the new imaging domain. The user can therefore obtain a more precise analysis, manually annotating GUVs in Label Studio, exporting the resulting JSON file and running the code GUV_analyser.py[43].

Once the corrected annotations are exported, the refined dataset can be used to retrain the model using the scripts provided in the repository *Training Section* [43], offering the possibility to train the model using RGB or grayscale images. This pipeline closes the loop between deployment and model refinement, allowing *VesiScope* to progressively adapt to new microscopes, sample types, or experimental conditions with minimal annotation overhead.

## Discussion

Quantitative characterization of GUV preparations commonly relies on fluorescent probes, which are essential for high-throughput analysis in flow cytometry or fluorescence microscopy. By enabling selective detection in the first case and enhancing contrast-to-noise ratio in the second, fluorescent labeling greatly facilitates quantitative measurements of vesicle yield and size distributions through automatic GUV detection. However, in many applications the fluorescent probe is not required for the intended experiment and is introduced solely for characterization purposes. Consequently, the analyzed sample differs from the target formulation, potentially leading to unaccounted alterations in vesicle yield, size distribution, membrane properties, or functional behavior. This limitation may impact the evaluation of GUV formulations across a wide range of applications. In membrane biophysics studies, for example, specific lipid compositions are used to assess membrane properties. Likewise, in transmembrane protein investigations, vesicle populations must be characterized without additional perturbations to assess the effect of protein incorporation. In synthetic biology or microrobotic applications, encapsulated cell-free expression systems, biochemical machineries, or nano-/micromotors may influence vesicle formation and stability. In all these cases, reliable quantification of GUV yield and size distribution is most informative when performed under the same label-free conditions used in downstream experiments.

Although phase-contrast microscopy enables label-free imaging, automated GUV detection remains challenging for traditional image processing pipelines, due to overlapping intensity distributions and low CNR[30]. In contrast, DL-based detection frameworks are inherently robust against these limitations and can accurately identify objects even under minimal contrast conditions, but existing AI-based methods still require fluorescent labelling[35,36], limiting their applicability to formulations that tolerate exogenous probes, and leaving unlabelled GUVs without a dedicated detection tool. To fill this gap, we introduced a DL approach for the automated detection and size characterization of GUVs in phase-contrast microscopy images.

To maximize accessibility, the trained model is distributed as an easy-to-install desktop application featuring an intuitive user interface and requiring neither programming expertise nor external dependencies or dedicated GPU hardware, following lightweight local deployment principles that ensure full reproducibility across different research environments. Its intuitive interface significantly reduces data analysis time by allowing the high-throughput processing of large datasets. Furthermore, the confidence threshold adjustment and the Label Studio-compatible JSON output embed human-in-the-loop correction and continuous training capabilities directly into the analysis workflow, allowing users to iteratively refine the model on new imaging conditions (or at least evaluate its performances) without the need of *de novo* annotations.

The model performance results demonstrate good generalization while supporting the use of lightweight architectures compatible with standard hardware. The substantial improvement of the grayscale model over the RGB model on the OOD dataset (Fig 2 C,D and S1 Table) supports the hypothesis that color information provides acquisition-specific features rather than biologically relevant information. Grayscale training therefore acts as an implicit domain-randomization strategy, reducing overfitting to illumination and sensor characteristics of the acquisition system, ultimately improving robustness across imaging conditions. At the same time, model complexity had minimal impact on performance, with the *nano* variant matching or outperforming larger models, demonstrating that accurate GUV detection can be achieved using computationally efficient architectures suitable for deployment on conventional laptops. The reduced detection performance for GUVs smaller than 5 µm can be attributed to their limited apparent contrast close to the optical resolution limit, which makes them intrinsically difficult to detect but also to manually annotate, resulting in a reduced number of such annotations in the training set (S1 Fig). In practice, this limitation can be mitigated by increasing microscope magnification, but this approach comes at the cost of longer acquisition times due to the reduced field of view. Therefore, the choice of magnification depends on the expected GUV size distribution, balancing throughput for larger vesicles against improved detection of smaller ones.

In conclusion, we present a stand-alone, user-friendly application for the automated detection and quantitative characterization of GUVs from phase-contrast microscopy images, complemented by a human-in-the-loop refinement pipeline(Fig 1). By automatically identifying label-free vesicles, VesiScope enables direct quantification of GUV yield and size distributions across a wide range of formulations and preparation methods without requiring the addition of fluorescent probes. This capability allows researchers to characterize GUV populations under the same conditions used for their intended applications, avoiding potential perturbations associated with fluorescent labeling. Although VesiScope is not designed to directly measure specific phenomena under investigation, it provides a robust and accessible framework for the quantitative characterization of the underlying GUV populations. By delivering rapid measurements of vesicle yield and size distribution, the software enables researchers to evaluate the quality and reproducibility of their GUV preparation protocols and to identify formulation or production issues before proceeding to downstream experiments. As a result, VesiScope can support a broad range of GUV-based applications, from membrane biophysics and synthetic biology to emerging microrobotic systems, by ensuring that downstream experiments are conducted on quantitatively characterized and reproducible vesicle populations.

**Fig 1:**
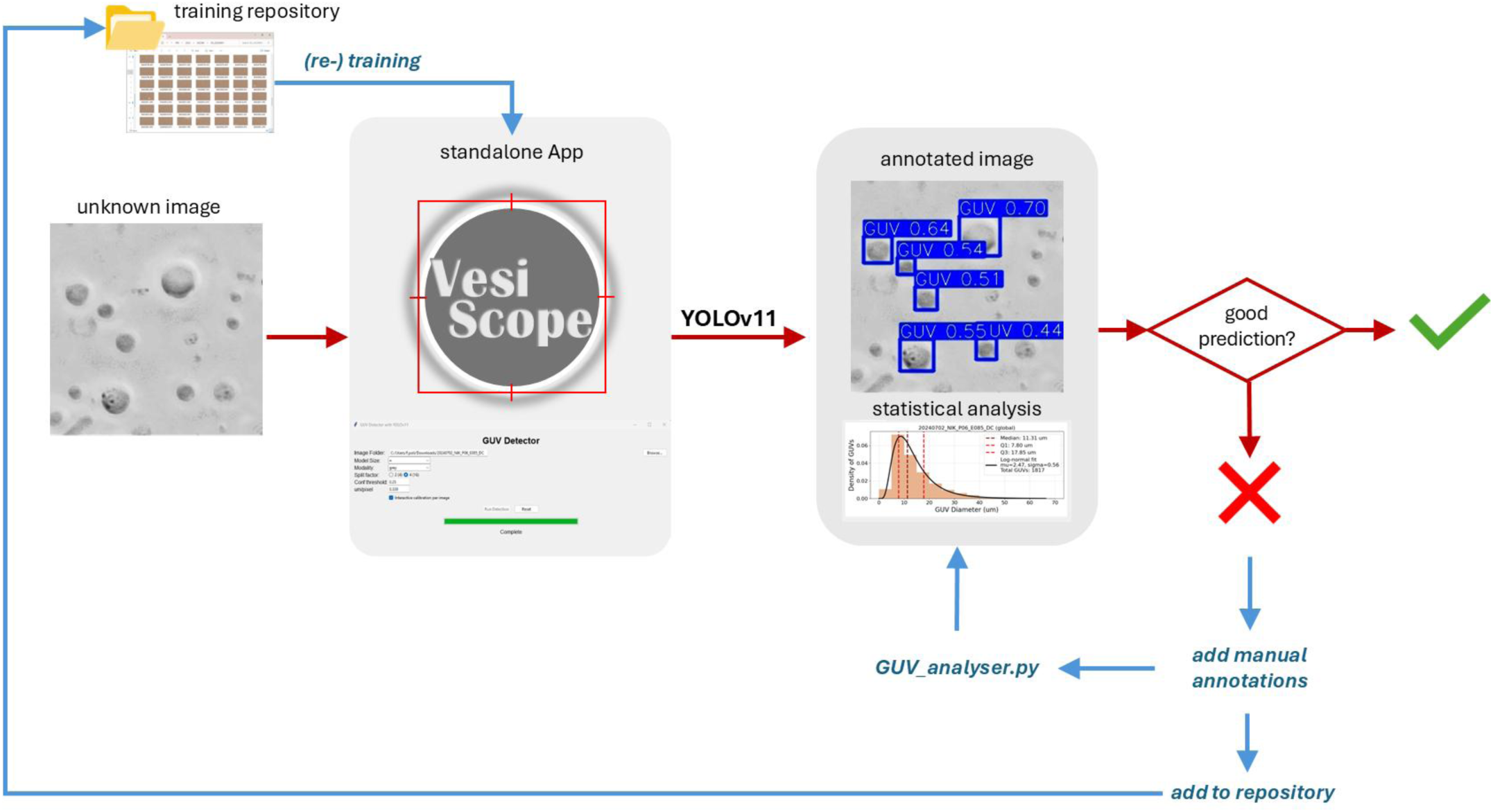
Human-in-the-loop pipeline for the quantification and size distribution analysis of GUV populations from phase-contrast microscopy images. Phase-contrast images are first processed by VesiScope, which uses a YOLOv11-based object detection model to automatically identify GUVs and generate quantitative outputs, including vesicle counts and size distributions. Predicted detections can then be reviewed by the user and, when necessary, exported to Label Studio for manual annotation and correction. The resulting annotations can be analyzed using the provided GUV_analyser.py script to obtain updated statistical measurements. Corrected annotations can subsequently be added to the training repository and used either to retrain the existing model or to train a new model from scratch. To facilitate this process, VesiScope is complemented by a publicly available GitHub repository and Google Colab notebooks that enable model training and refinement on standard hardware. This human-in-the-loop workflow supports continuous model improvement and adaptation to different phase-contrast microscopes, imaging conditions, and GUV formulations while maintaining an accessible and reproducible analysis pipeline.

**Fig 2:**
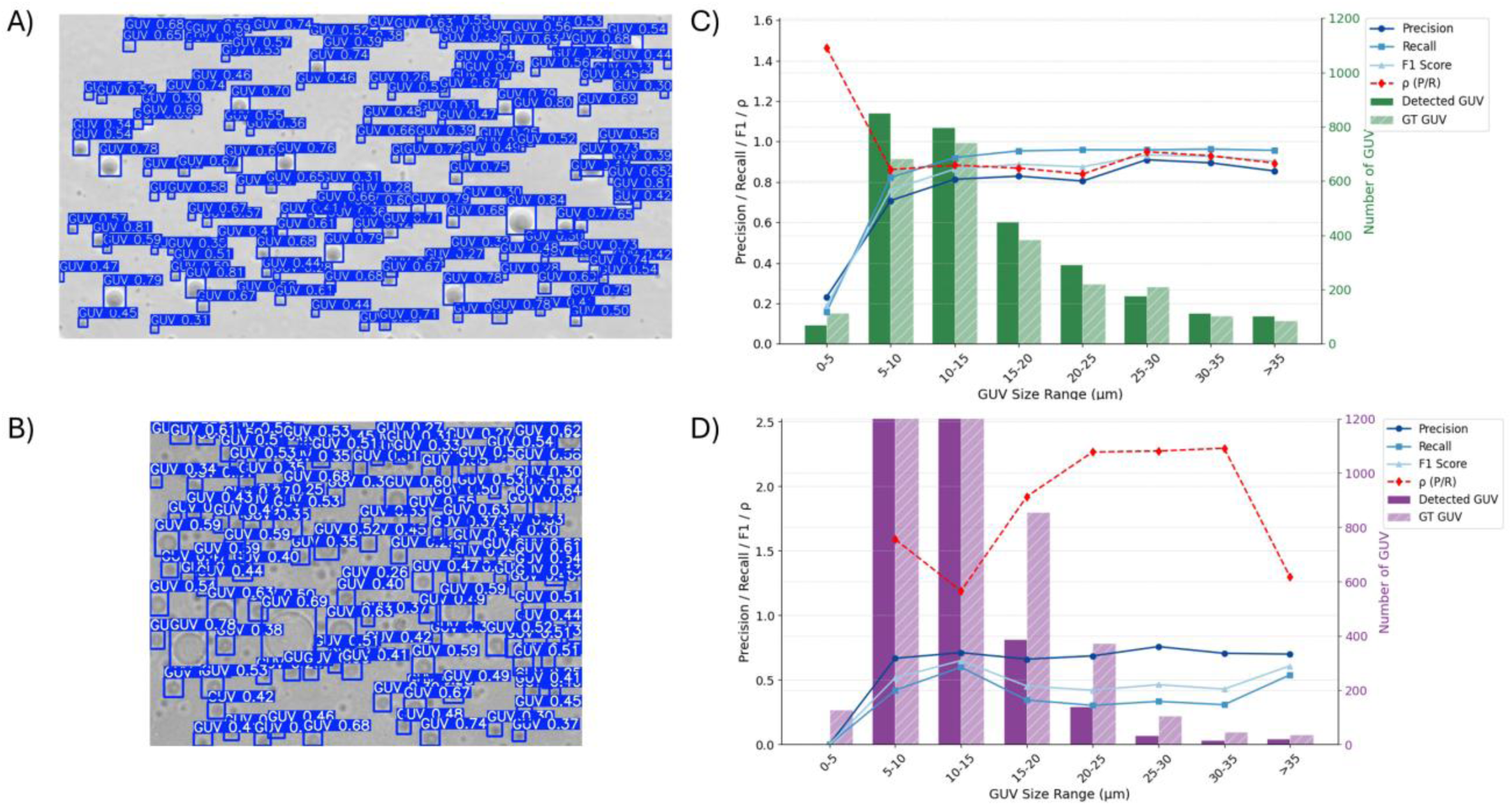
Performances on ID and OOD dataset of YOLOv11 nano model trained on grayscale images. A) and B) With a confidence threshold = 0.25 and IoU = 0.50, the model is able to detect the majority of GUVs both in ID (A) and OOD (B) dataset as shown in the example images generated by VesiScope where GUVs are framed in bounding boxes with the associated confidence scores. C) and D) The comparison between manually annotated images and model predictions show similar values in ID dataset (C) for GT (dashed bar) and predicted (full bars) GUVs stratified by diameters in 5 µm bins, resulting in high values (above 0.8) for Precision (circles), Recall (squares), and F1 score (triangles) and a constant ρ value (diamonds) for GUVs above 5 µm in diameters. As expected, the model shows reduced but still good performances in OOD dataset (D) for GUVs with diameters above 5 µm, with lower Precision and Recall values (above 0.5) and more heterogenous ρ values.

**Fig 3:**
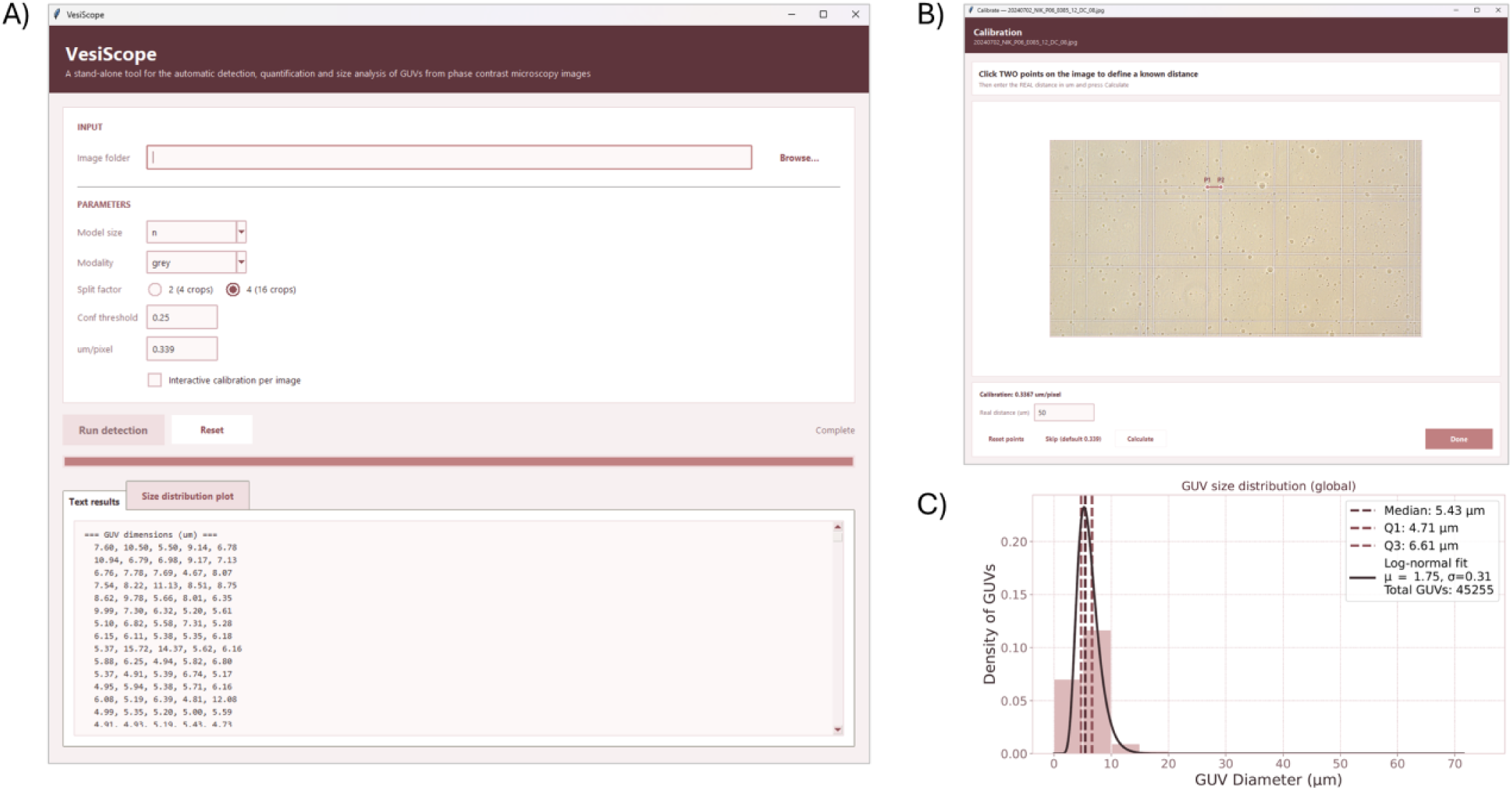
VesiScope user interface. A) The stand-alone desktop application allows single or batch processing of phase-contrast images both in original RGB format or through grayscale compression for improved generalization. B) When μm/pixel conversion factor is unknown, an interactive window allows the calculation through the selection of two points with known distance, like those from a scale bar or from a counting chamber. C) Size distributions can be easily visualized in a plot generated by the application, together with a CSV file containing the list of diameters expressed in μm for downstream analysis.

## Acknowledgements

We thank Elisa Roberti, Gaia Petrucci, Francesco Bianciardi and Jyoti Sharma for contributing to manual annotation of GUVs from phase contrast images.

## Supporting information

**S1 Fig.**
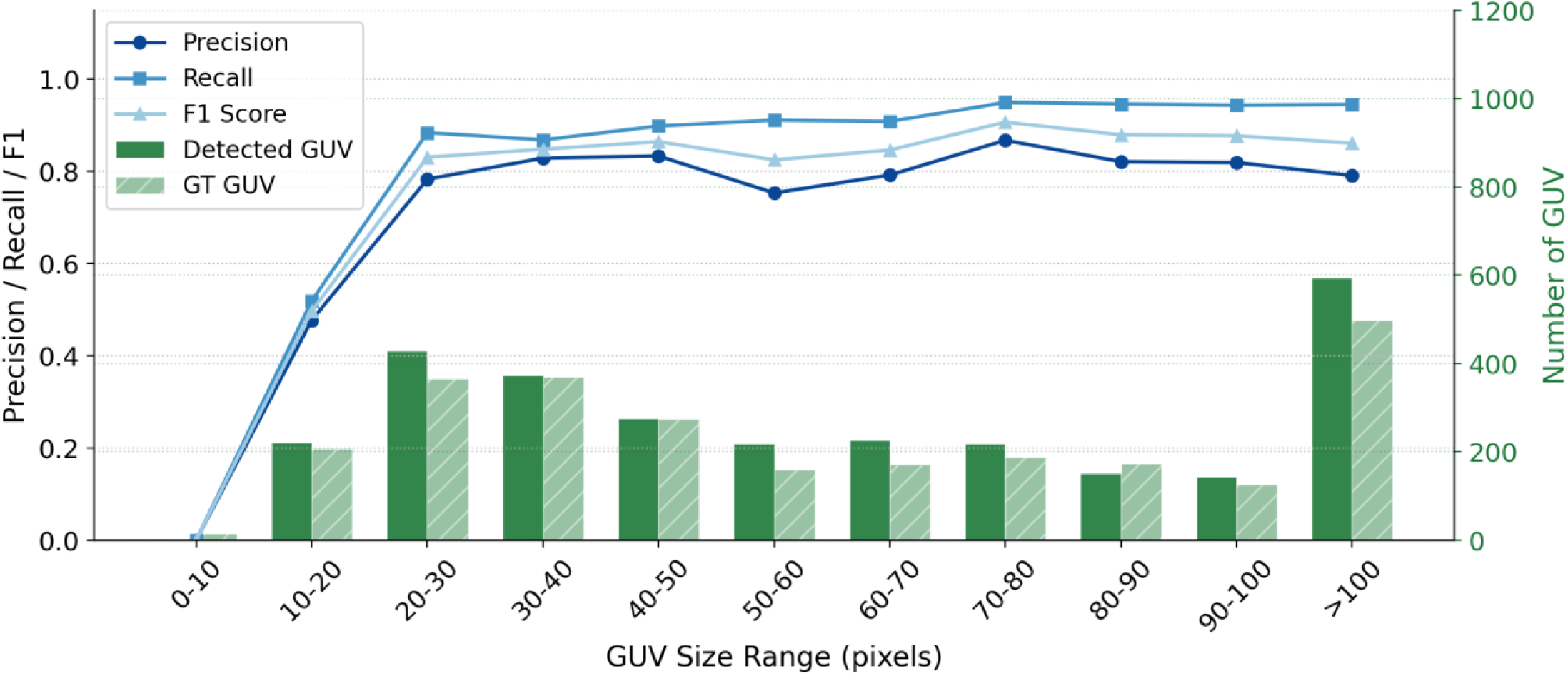
Precision, Recall, and F1 score of the greyscale YOLOv11-n model stratified by GUV diameter expressed in pixels on the ID test set. Bar heights (right axis) indicate the number of ground-truth (GT) and detected GUVs in each size bin. Detection performance plateaus with 20-pixels diameters, corresponding to ∼2.5% of the processed image size.

**S1 Table.** Detection performance (Precision, Recall, mAP50) for YOLOv11 nano (n), small (s), and medium (m) variants trained on RGB and greyscale images on both test set (in-distribution) and cross-microscope generalization set (out-of-distribution).

| **Dataset** | **Model** | **Params (M)** | **Precision (RGB)** | **Recall (RGB)** | **mAP50 (RGB)** | **Precision (Grey)** | **Recall (Grey)** | **mAP50 (Grey)** |
|---|---|---|---|---|---|---|---|---|
| **ID** | n | 2.5 | 0.845 | 0.844 | 0.875 | 0.868 | 0.842 | 0.881 |
| **ID** | s | 9 | 0.858 | 0.833 | 0.866 | 0.856 | 0.847 | 0.872 |
| **ID** | m | 20 | 0.867 | 0.815 | 0.863 | 0.853 | 0.832 | 0.874 |
| **OOD** | n | 2.5 | 0.319 | 0.105 | 0.198 | 0.603 | 0.517 | 0.514 |
| **OOD** | s | 9 | 0.205 | 0.115 | 0.137 | 0.643 | 0.532 | 0.525 |
| **OOD** | m | 20 | 0.157 | 0.200 | 0.144 | 0.666 | 0.459 | 0.519 |

